# VPS18 recruits VPS41 to the human HOPS complex via a RING-RING interaction

**DOI:** 10.1101/169185

**Authors:** Morag R. Hunter, Edward J. Scourfield, Edward Emmott, Stephen C. Graham

**Affiliations:** Department of Pathology, University of Cambridge, Cambridge, United Kingdom; Current affilitation: Faculty of Life Sciences & Medicine, Department of Infectious Diseases, King’s College London, London, United Kingdom

**Keywords:** Class C core, Sec1/Munc18, autophagy

## Abstract

Eukaryotic cells use conserved multisubunit membrane tethering complexes, including CORVET and HOPS, to control the fusion of endomembranes. These complexes have been extensively studied in yeast, but to date there have been far fewer studies of metazoan CORVET and HOPS. Both of these complexes comprise six subunits: a common four-subunit core and two unique subunits. Once assembled, these complexes function to recognise specific endosomal membrane markers and facilitate SNARE-mediated membrane fusion. CORVET promotes the homotypic fusion of early endosomes, while HOPS promotes the fusion of lysosomes to late endosomes and autophagosomes. Many of the subunits of both CORVET and HOPS contain putative C-terminal zinc-finger domains. Here, the contribution of these domains to the assembly of the human CORVET and HOPS complexes has been examined. Using biochemical techniques, we demonstrate that the zinc-containing RING domains of human VPS18 and VPS41 interact directly to form a stable heterodimer. In cells, these RING domains are able to integrate into endogenous HOPS, showing that the VPS18 RING domain is required to recruit VPS41 to the core complex subunits. Importantly, this mechanism is not conserved throughout eukaryotes, as yeast Vps41 does not contain a C-terminal zinc-finger motif. The subunit analogous to VPS41 in human CORVET is VPS8, in which the RING domain has an additional C-terminal segment that is predicted to be disordered. Both the RING and disordered C-terminal domains are required for integration of VPS8 into endogenous CORVET complexes, suggesting that HOPS and CORVET recruit VPS41 and VPS8 via distinct molecular interactions.

## INTRODUCTION

Eukaryotic cells exert tight control over the fusion of their membrane-bound compartments. SNARE-mediated membrane fusion events are regulated by multiple proteins (1, 2), including multisubunit membrane tethering complexes. The class C core vacuole/endosome tethering (CORVET) and homotypic fusion and vacuole protein sorting (HOPS) complexes mediate homotypic fusion of early endosomes, and heterotypic fusion of late endosomes to vacuoles (in yeast) or lysosomes (in higher eukaryotes), respectively (figure 1) (3-5). HOPS is also required for the fusion of autophagosomes to lysosomes (6–9). In mammalian cells, these complexes share four core subunits (VPS11, VPS16, VPS33A and VPS18, collectively known as the class C core), along with two subunits unique for CORVET (VPS8, TRAP-1) or HOPS (VPS39, VPS41) (figure 1) (5, 10, 11). These non-core subunits direct the complexes to their different membrane targets: Rab5-positive endosomes for CORVET (5, 12), and Rab7-positive for HOPS (13, 14).

**Figure 1:**
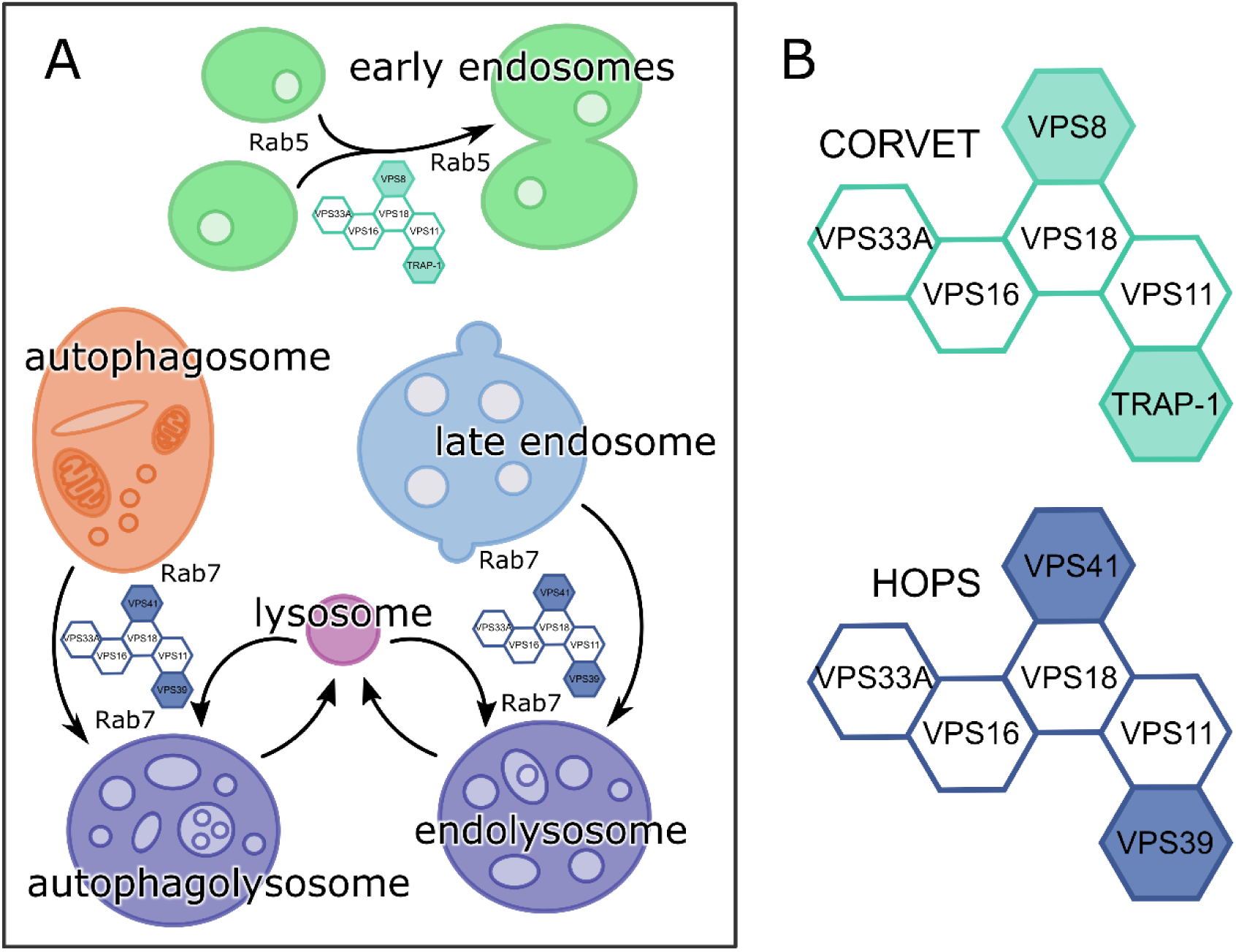
CORVET and HOPS facilitate SNARE-mediated membrane fusion events in the endo-lysosomal and autophagosomal systems. (A) CORVET mediates fusion of Rab5-positive membranes (early endosomes) and HOPS mediates fusion of the Rab7-positive membranes (late endosomes to lysosomes, autophagosomes to lysosomes). (B) CORVET and HOPS share a similar architecture, with a common four-subunit core and two unique subunits each.

Two electron microscopy structures have been obtained for yeast HOPS (15, 16), showing the general structure of this complex. Based on the first HOPS structure (stabilised by glutaraldehyde cross-linking), the complex has been described as seahorse-shaped, with a “head” and “tail” formed of globular subunits (15). More recently, a second, non-cross-linked, structure has been published, showing HOPS to have long, flexible, extended legs (16). While obtained under different conditions, both of these studies concluded that Vps41 contacts the class C core subunits at or near Vps18. The Vps18-Vps41 contact site is also approximately where the Vps16-Vps33 subcomplex and Vps11 join the larger complex (the “head”). Meanwhile, Vps39 appears to only contact Vps11 at the opposite (“tail”) end of the complex.

To date, the structure of mammalian HOPS has been assumed to be the same as that of the yeast HOPS complex. However, there is evidence of evolutionary divergence within the membrane trafficking system. Metazoan VPS33A and VPS33B are both homologs of the yeast HOPS subunit Vps33 (17), but only VPS33A is found in human HOPS (6, 11, 18) and mutations in these two homologs generate significantly different phenotypes (19–23). Also, Vps41 in yeast has been shown to bind directly to Ypt7 (24), but in humans it appears that the analogous interaction (with Rab7) is mediated by an additional protein, RILP (14). There is also evidence that HOPS can interact with Arl8b (a small GTPase) on lysosomal membranes (25, 26). Furthermore, there is limited direct biochemical data pertaining to the organisation of the human HOPS complex and its interaction with membrane markers. Accordingly, we set out to characterise the functional domains of the human HOPS subunits.

Aside from the Sec1/Munc18-like component VPS33A (18), the subunits of CORVET and HOPS have a very similar architecture: a β-propeller domain at the N terminus (27), followed by an α-solenoid and (usually) a C-terminal zinc-finger domain (28–31). For human VPS11, VPS18 and VPS41, these C-terminal zinc finger domains match the consensus sequence of the really interesting new gene (RING) domain (31, 32). The C terminus of yeast HOPS subunits have been broadly implicated in complex assembly (e.g. (33)), but without special attention paid to the RING domains. Further, it should be noted that no zinc-finger motif is present in yeast Vps41(31). RING domains are a type of zinc finger, with a motif consisting of eight residues (six or seven cysteines and one or two histidines) arranged to bind two Zn^2+^ ions, and forming a “cross brace” structure (34, 35). As RING domains can mediate protein-protein interactions (36, 37) or self-assemble (38), we aimed to characterise the role of these domains in human HOPS subunits. While we initially hypothesised that these RING domains may mediate recognition of membrane markers by HOPS, this was not supported by our preliminary data. Instead, the RING domains we investigated were found to mediate the association of the HOPS subunits within the complex itself.

## RESULTS

The boundaries of the RING-H2 domains (with the zinc ligands ordered as C3H2C3) of HOPS subunits VPS18 and VPS41 were defined by analysing predicted HOPS protein secondary structure using NETSURFP (39) and homology to RING domains of known structure using HHPRED (40). The RING-H2 domain of VPS18 contains an extended space between the sixth and seventh zinc ligands (65 amino acid residues, rather than the typical maximum of 48 (35, 41)), while otherwise maintaining the characteristic spacing of the zinc ligands. This would be expected to result in a longer disordered loop, but otherwise maintain all other characteristics of this zinc-binding motif. The putative C-terminal zinc finger domain of VPS39 is significantly shorter than those of VPS18 and VPS41, containing only four Zn^2+^ ligands, and the closest homologue of known structure is the zinc finger domain of yeast Pcf11 (42).

A yeast two-hybrid screen was performed to identify potential interaction partners of the human VPS18 and VPS41 RING domains and the VPS39 zinc finger domain (supplementary figure 1 and supplementary data). We hypothesised that these domains may identify markers of membrane identity (e.g. Rab proteins), based on evidence that RING domains may mediate recruitment of mBCA2 (a.k.a. Rabring7, (37)) and hVPS41 (43) to membranes. However, attempts to demonstrate direct binding between these domains and selected proteins identified in the yeast two-hybrid screen were unsuccessful, suggesting that these putative interactions were ‘false positives’. We thus hypothesised that, rather than binding to other cellular factors, the VPS18 and VPS41 RING domains may instead mediate intra-molecular interactions within the HOPS complex.

This hypothesis was tested by performing pulldown experiments using recombinant proteins. The RING domains of VPS18 (residues 842–973) and VPS41 (residues 778–854) were expressed as GST fusion proteins and purified by glutathione immobilisation affinity chromatography followed by size-exclusion chromatography (SEC). All six subunits of the human HOPS complex were expressed individually with N-terminal myc epitope tags using a eukaryotic (wheat germ) in vitro transcription/translation expression system. Affinity capture (pulldown) experiments were performed by incubating the cell-free expression lysates with GST-tagged VPS18 RING or VPS41 RING domains immobilised on glutathione resin (figure 2A). These experiments showed that the VPS18 RING construct interacted exclusively with full length VPS41, and that VPS41 RING interacted exclusively with full length VPS18.

**Figure 2:**
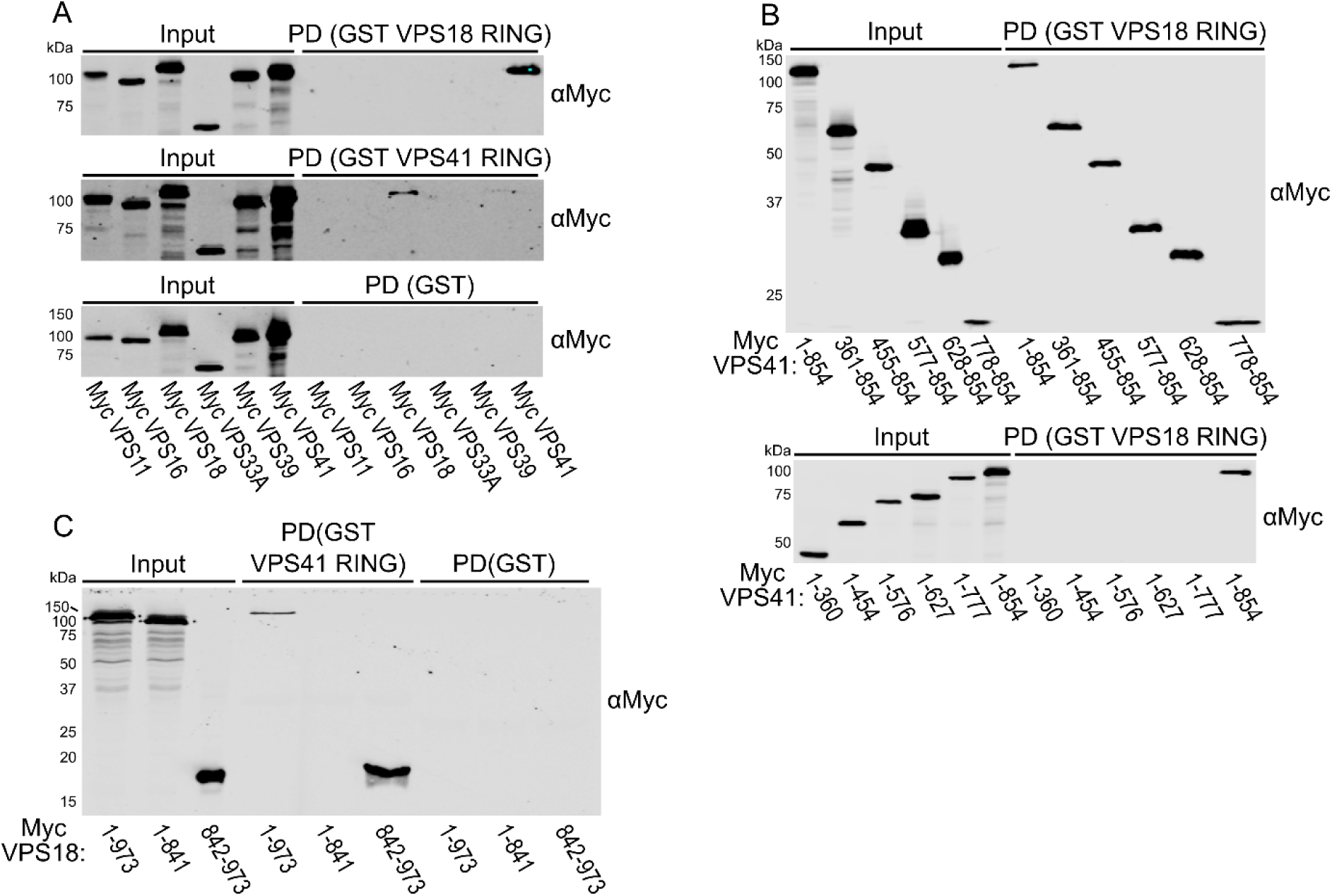
The RING domains of VPS18 and VPS41 are sufficient to bind to full length HOPS subunits. (A) Myc-tagged HOPS subunits produced by in vitro transcription/translation were subjected to GST pulldown (PD) using GST-tagged VPS18 and VPS41 RING domains or GST alone as bait proteins followed by immunoblotting (anti-myc). (B & C) A series of myc-tagged VPS41 and VPS18 truncations were produced by in vitro transcription/translation, then subjected to GST pulldown with the indicated bait proteins followed by immunoblotting (anti-myc).

A series of VPS41 truncations were made to delineate the regions necessary for binding the VPS18 RING domain. This showed that only VPS41 constructs including the C-terminal RING domain (residues 778-854) were able to interact with the VPS18 RING domain (figure 2B). Analogously, full length VPS18 was compared to VPS18 constructs with the C-terminal RING domain removed (residues 1-841) or encoding just the RING domain alone (residues 842-973). This experiment showed that the RING domain of VPS18 is necessary and sufficient for binding to the VPS41 RING domain in vitro (figure 2C). Accordingly, we proceeded to directly assess the interaction between purified VPS18 and VPS41 RING domains.

Reciprocal GST pulldown experiments using all purified recombinant components showed that VPS18 and VPS41 RING domains form a direct interaction (figure 3A). His-tagged VPS18 RING was purified by Ni^2+^ immobilisation affinity chromatography followed by SEC, while tag-free VPS41 RING was purified by removal of the GST tag from GST-VPS41 RING using human rhinovirus 3C protease and subsequent glutathione capture then SEC. The intensity of Coomassie-stained SDS-PAGE bands from the GST pulldown experiment suggested that the VPS18 and VPS41 RING domains form an equimolar complex. To characterise the stoichiometry of the complex directly, the mass of the complex was determined using SEC with inline multiangle light scattering (MALS). SEC-MALS analysis showed both untagged VPS41 RING and His-tagged VPS18 RING to be monomeric in solution with average molecular masses of 9.5 kDa and 15.6 kDa, respectively (figure 3B). The VPS18H_6_ + VPS41 RING domain complex co-eluted during the SEC-MALS experiment (figure 3C), the molecular mass of the complex (22.8 kDa) being consistent with these proteins forming a 1:1 heterodimer (figure 3B).

**Figure 3:**
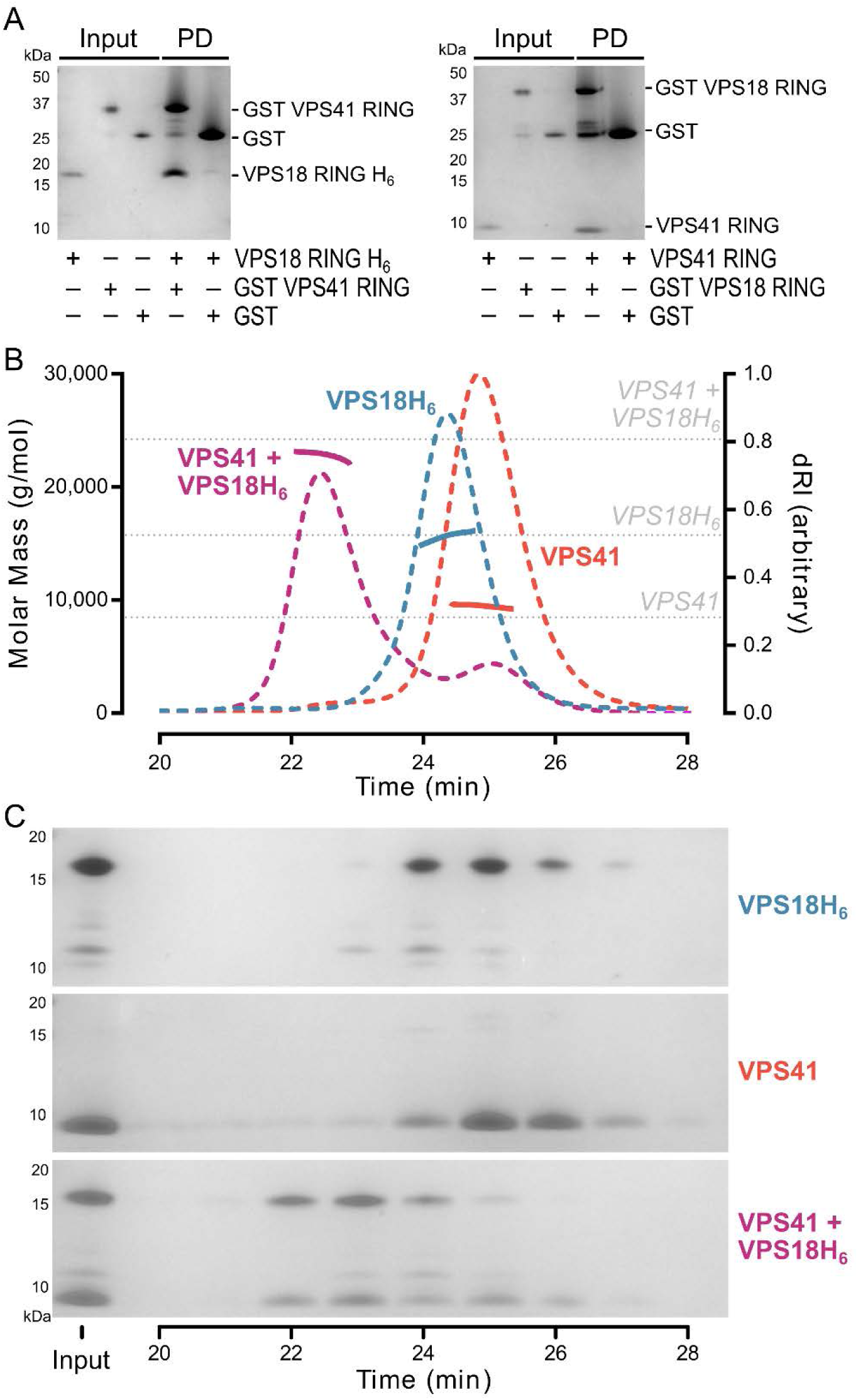
The RING domains of VPS41 and VPS18 form a direct heterodimer. (A) Purified recombinant VPS41 and VPS18 RING domains were subjected to GST pulldown (PD) experiments using the indicated bait proteins followed by SDS-PAGE (Coomassie). (B) SEC-MALS of VPS18 and VPS41 RING domains alone or in complex. Weight-averaged molar masses (solid lines) are shown across the elution profiles (differential refractive index, dashed lines). The expected molar masses of the monomeric individual RING domains and of a 1:1 heterodimeric complex are also shown (grey dotted lines). (C) SDS-PAGE of fractions from SEC-MALS experiments shown in (B), confirming that VPS41 and His-tagged VPS18 RING domains co-elute from the SEC-MALS experiment as a complex.

Having demonstrated that the RING domains of VPS18 and VPS41 RING form a direct, stable heterodimeric complex in vitro, we sought to probe the relevance of this interaction for HOPS assembly in human cells. HEK293T cells were transfected with EGFP-tagged VPS18 and VPS41 constructs (full length and various truncations, figure 4A). Immunoprecipitations were performed to assess whether these transfected constructs could interact with the HOPS complex. We assessed the interactions of our transfected EGFP-tagged constructs solely with endogenously-expressed HOPS subunits rather than co-transfecting multiple subunits, so as to reduce the risk of false-positive interactions which can occur through co-overexpression (44).

**Figure 4:**
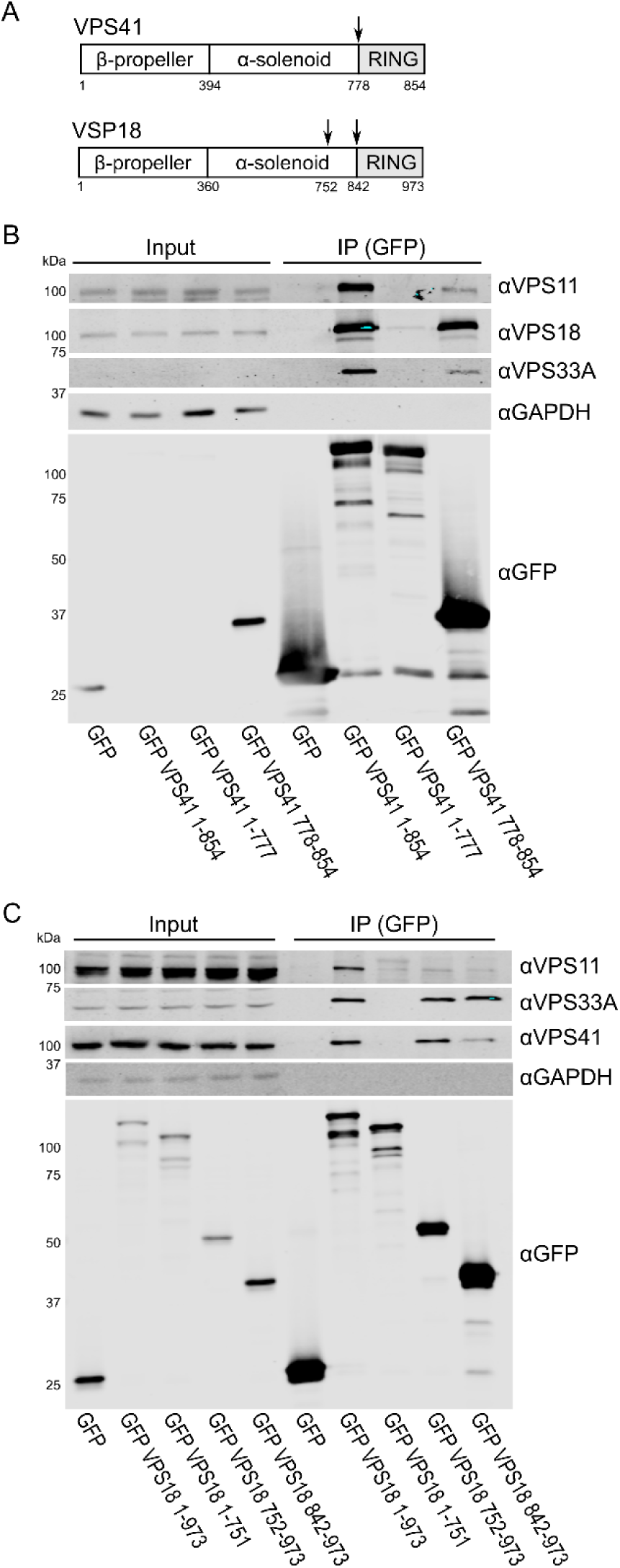
The RING domains of VPS41 and VPS18 are required for efficient interaction with endogenous HOPS components. HEK293T cells were transfected with GFP-tagged VPS41 and VPS18 truncation constructs. After immunoprecipitation (IP), samples were immunoblotted for the indicated endogenous HOPS subunits (VPS11, VPS18, VPS33A, and/or VPS41), GAPDH (loading control), and GFP. (A) Schematic showing truncation constructs. (B) The RING domain of VPS41 (778-854) integrates into the endogenous HOPS complex. (C) While the RING domain of VPS18 (842-973) is able to interact with the endogenous HOPS complex, an extended construct (752-973) is required to co-immunoprecipitate all HOPS subunits as efficiently as full length (1-973) VPS18.

When various VPS41 constructs were transfected into HEK293T cells, the results were in agreement with those of the pulldown experiments: the VPS41 RING domain (residues 778–854) is sufficient and necessary to co-immunoprecipitate endogenous VPS18, and does so with an efficiency equivalent to that of full length VPS41 (figure 4B). The VPS41 RING domain was, however, only able to co-immunoprecipitate a fraction of the endogenous VPS11 and VPS33A compared to the full length construct, perhaps due to steric hindrance of the GFP tag, or because the longer VPS41 contributes to a more stable configuration of the HOPS complex. The complementary N-terminal construct (residues 1–777) did not interact with either VPS11 or VPS33A, indicating that the RING domain is absolutely required for recruitment of VPS41 to the HOPS complex.

Next, we transfected HEK293T cells with the VPS18 RING construct (residues 842–973), While this construct was sufficient to bind endogenous VPS41, it was not as efficient in co-immunoprecipitating VPS41 as the full-length VPS18 construct (figure 4C, far right lane). We also observed that the complementary N-terminal VPS18 construct (residues 1–841) was able to co-immunoprecipitate a small amount of endogenous VPS41 (supplementary figure 2A). Thus, we performed a series of N- and C-terminal truncations of VPS18 and tested these for their ability to co-immunoprecipite endogenous HOPS components in transfected cells. This experiment showed that a portion of the VPS18 α-solenoid contributes to binding of VPS41 in cells, with VPS18 residues 752–973 comprising the entire VPS41 binding region (figure 4C, supplementary figure 2B). These results are broadly in agreement with our in vitro results, showing that the VPS18 RING domain is sufficient to bind VPS41. The lack of binding between VPS18 residues 1–841 and VPS41 in biochemical solution may arise from an inability of this region of VPS18 to fold correctly in the absence of other HOPS components.

In addition to binding VPS41, all VPS18 constructs containing the RING domain (residues 842-942) co-immunoprecipitate endogenous VPS33A to approximately the same extent as full-length VPS18. This suggests that the RING domain itself is sufficient to mediate this interaction and that the interaction is unaffected by the recruitment of VPS41 (figure 4C). As VPS16 is required for recruitment of VPS33A to HOPS (6, 18), we can infer that VPS16 is also recruited to HOPS by the VPS18 RING domain. Furthermore, only constructs of VPS18 which contained the β-propeller and a large N-terminal portion of the α-solenoid were able to co-immunoprecipitate endogenous VPS11 (supplementary figure 2C), consistent with previous observations in yeast (27). The VPS18 RING domain does not appear to contribute to binding of VPS11 to HOPS (supplementary figure 2C).

VPS41 in HOPS is analogous to VPS8 in CORVET, while VPS18 is common to both complexes (figure 1) (4, 5). As with VPS41, VPS8 also contains a RING domain, but it has an additional region C-terminal to the RING domain that is predicted to be disordered (figure 5A, B). We investigated whether the VPS8 RING domain interacts with the class C core of CORVET by an equivalent mechanism to VPS41 in HOPS. The RING domain of VPS8 was expressed in E. coli as a GST fusion either with or without the C-terminal putative disordered region (VPS8 RING long and VPS8 RING short, respectively). Surprisingly, neither GST VPS8 RING long nor GST VPS8 RING short interacted with His-tagged VPS18 RING in pulldown experiments (figure 5C). To probe whether a region of VPS8 N-terminal to the RING domain was required for the interaction, pulldown experiments were performed to assess the binding of full length VPS8 or VPS41 (expressed by in vitro transcription/translation) to purified recombinant VPS18 RING domain. Again, we found that VPS8 did not bind the purified VPS18 RING domain (figure 5D).

**Figure 5:**
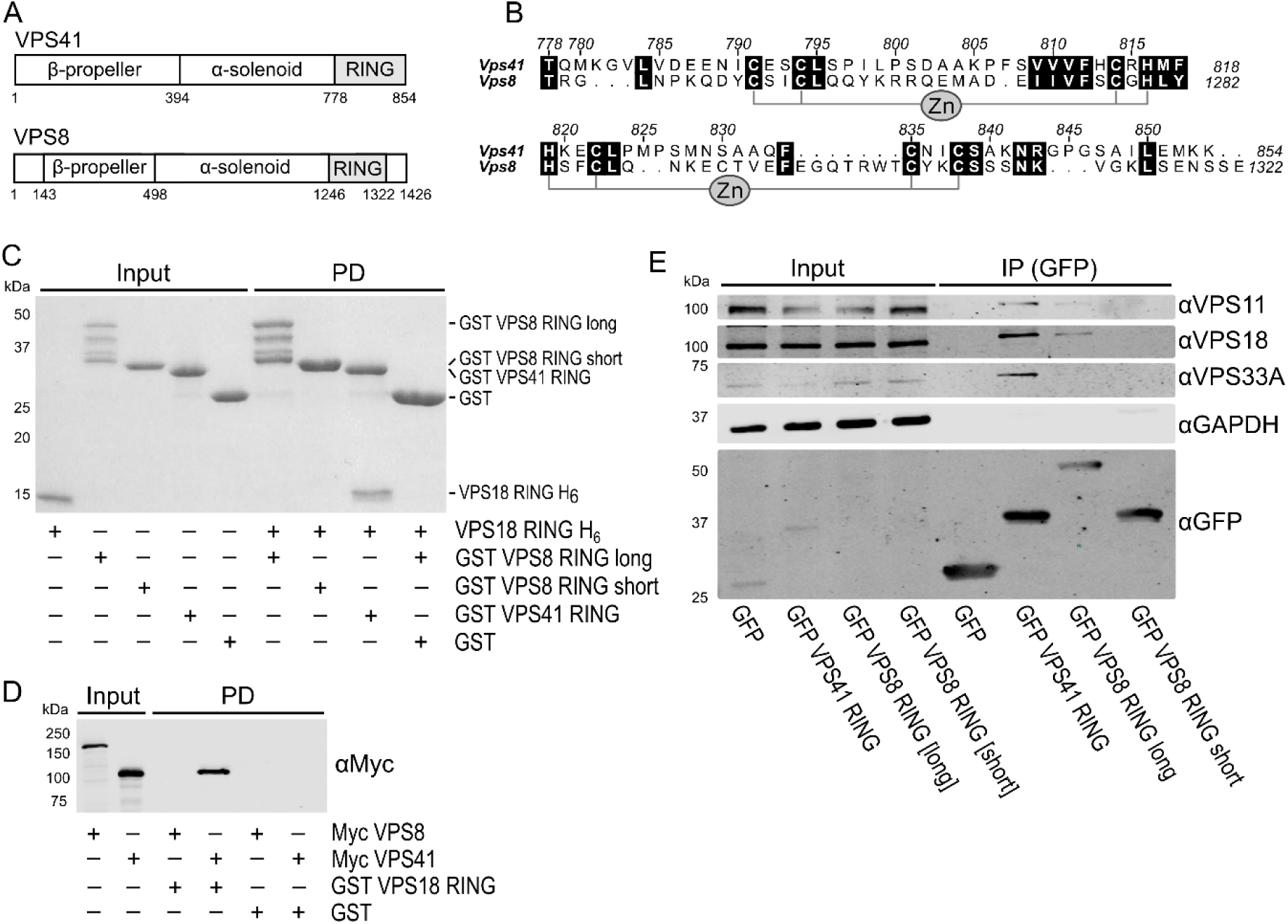
VPS41 and VPS8 share a similar domain structure, but the VPS8 RING domain is not sufficient to bind to VPS18. (A) Schematic representation of VPS41 and VPS8. (B) Schematic representation of the RING domain homology between VPS41 and VPS8. (C) GST pulldown (PD) experiments show that purified VPS18 RING domain is efficiently captured by the VPS41 RING domain, but not by the VPS8 RING domain. (D) Full length VPS41 produced by in vitro transcription/translation is pulled down by VPS18 RING domain, but full length VPS8 is not. (E) In HEK293T cells, the VPS8 RING long domain co-immunoprecipitates endogenous VPS18 – albeit less efficiently than the VPS41 RING domain – but the VPS8 RING short construct does not.

A lack of observable interaction between VPS8 and purified VPS18 RING domain could arise due to incorrect folding of the VPS8 domain when expressed recombinantly, due to a lack of required post-translational modifications, or due to a genuine lack of interaction between these domains. To discount the former two hypotheses, we tested the ability of VPS8 RING domain constructs to bind endogenous CORVET components in HEK293T cells. In cells, the VPS8 RING long construct (containing both the RING domain and a disordered C terminus) did appear to be able to integrate with endogenous CORVET complexes. It was able to co-immunoprecipitate endogenous VPS18 and VPS11 from these cells, albeit with considerably lower efficiency than VPS41 RING (figure 5E). This is in line with the previous finding that yeast Vps8 requires only its C terminus (broadly defined as Vps8 residue 450 (in the α-solenoid) to the end) to be recruited to CORVET (45). Co-immunoprecipitation with endogenous VPS33A was not detected, although it may have been below the level of detection afforded by this antibody. The VPS8 RING long construct had a lower co-immunoprecipitation efficiency for all core subunits compared to VPS41 RING, which may be due to either the relative abundance or stability of HOPS versus CORVET in this cell line. Unfortunately, there were no commercially-available antibodies raised against VPS8 which could detect endogenous protein by western blot in HEK293T cells (see ‘Materials and Methods: Antibodies’), so we were unable to test whether endogenous VPS8 interacts with VPS18 truncation constructs.

## DISCUSSION

We sought to assess the contribution of the zinc-containing domains of VPS18, VPS39 and VPS41 to the function of the mammalian HOPS complex. We showed using purified recombinant proteins that the RING domains of VPS18 and VPS41 form a stable, direct heterodimer and that this interaction mediates recruitment of VPS41 to the HOPS complex in mammalian cells.

Our finding that the VPS18 and VPS41 RING domains mediate HOPS assembly is in broad agreement with a previous study in which co-transfected, epitope-tagged HOPS subunits were co-immunoprecipitated by truncated forms of VPS18 (containing the RING domain) and VPS41 (containing α-solenoid and RING) (11). Consistent with this study, our results show that the RING domain of VPS18 mediates interactions with both VPS41 and VPS16/VPS33A, the portion of HOPS described as the “head” of a seahorse-shaped complex (15), and we have refined the precise residue ranges required for binding of VPS41 and VPS18 to other HOPS subunits. While it was not possible to compare the efficiency of co-immunoprecipitation between truncation constructs in the previous study, our results show that minimal binding domains (VPS41 778–854 and VPS18 752–973) bind endogenous HOPS components with the same efficiency as the full-length proteins. Additionally, we have confirmed that VPS11 joins the HOPS “head” through contact with the VPS18 β-propeller and α-solenoid, and that this interaction does not require the VPS18 RING domain.

Unlike VPS41, the isolated RING domain of VPS8 was not sufficient to maintain an interaction with VPS18. In co-immunoprecipitation experiments the VPS8 RING short construct (containing only the RING domain, and therefore most similar to the VPS41 RING construct) did not demonstrate observable interactions with any endogenous HOPS/CORVET subunit we tested. This suggests that there is a qualitative difference between the class C core binding mechanism of VPS41 and VPS8; more than the RING domain of VPS8 is required for its recruitment to CORVET. Although the VPS8 sequence generally conforms to the RING-H2 domain consensus, it also contains a tryptophan residue two residues before the seventh zinc ligand — a characteristic feature of the closely-related plant homeodomain (PHD) motif (41, 46). In PHD domains, this tryptophan residue contributes to the hydrophobic core and influences the orientation of the C terminus of this motif (46, 47). This may explain the requirement of the VPS8 extended C terminus for binding to VPS18, although a complete understanding of how the VPS18 RING binds VPS41 and VPS8 awaits high-resolution structural characterisation.

To date, the configuration of mammalian HOPS has largely been inferred from studies of the homologous yeast complex. Here we demonstrate that the VPS41 RING domain is required for recruitment of VPS41 to the human HOPS complex. However, yeast Vps41 does not contain any zinc-finger motifs (31). Our findings thus indicate that the molecular interfaces that mediate the assembly of mammalian and yeast HOPS complexes are not identical. Based on the results presented in this study, a model of human HOPS assembly (figure 6) can be summarised as follows: VPS18 is central to HOPS assembly; The RING domain of VPS18 forms a hub that recruits the RING domain of VPS41 to HOPS; The VPS16-VPS33A subcomplex also connects to this hub via VPS16 (6, 18); VPS11, with VPS39 attached to its distal end (15), joins HOPS through contacts across an extended portion of the VPS18 N terminus. Overall, our data demonstrate that the interactions of RING domains are integral to the assembly of the mammalian HOPS complex, and that the molecular architectures of HOPS differ between yeast and humans.

**Figure 6:**
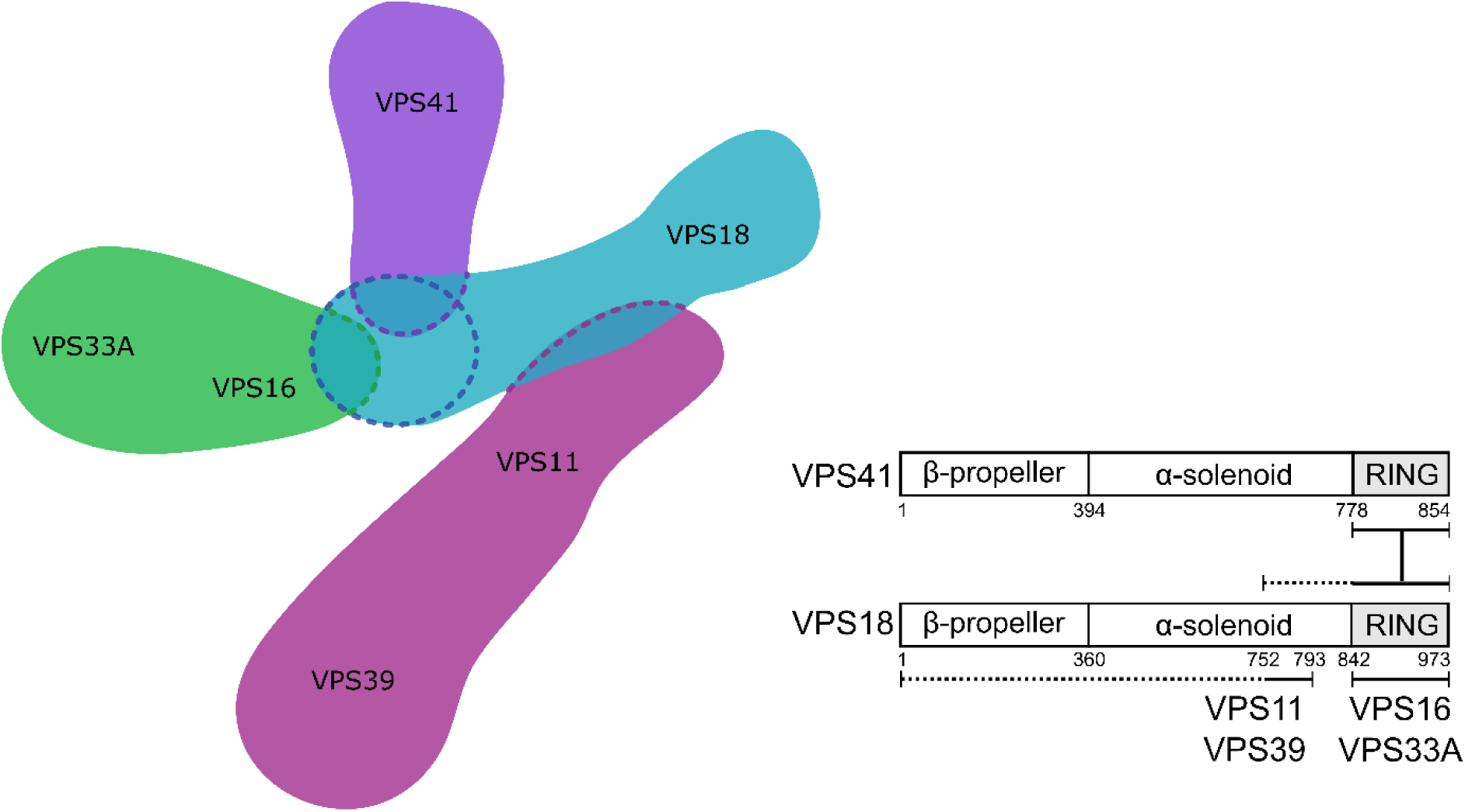
Model of HOPS assembly. The VPS18 RING domain acts as a hub for HOPS assembly, recruiting VPS41 (via its RING domain) and the VPS16-VPS33A subcomplex. The N terminus of VPS18 mediates interaction with the VPS11-VPS39 subcomplex, with residues 752-793 being essential.

## MATERIALS AND METHODS

### Antibodies

The following primary antibodies were used for immunoblotting: monoclonal anti-myc (Millipore, cat. 05-724); polyclonal anti-VPS11 (Proteintech, cat. 19140-1-AP, lot 44-161-2324577); polyclonal anti-VPS18 and polyclonal anti-VPS33A (as described in (18)); monoclonal anti-VPS41 (Santa Cruz, cat. sc-377118); monoclonal anti-GAPDH (Life Technologies, cat. AM4300); polyclonal anti-GFP (Sigma, cat. G1544). Three antibodies against VPS8 were tested but failed our validation assays for immunoblotting: polyclonal anti-VPS8 (Proteintech, cat. 15079-1-AP), polyclonal anti-VPS8 (Sigma, cat. HPA036871, lot R33997), and monoclonal anti-VPS8 (Sigma, cat. WH0023355M1-100UG). Antibodies were validated for western blotting by testing samples of protein from cell-free expression reactions, whole-cell lysates (containing transfected EGFP-tagged protein and/or endogenous protein), and immunoprecipitated samples from whole-cell lysates. IRDye 800CW-conjugated secondary antibodies for immunoblotting were supplied by LI-COR: goat anti-mouse (cat. 925-32210), goat anti-rabbit (cat. 925-32211), and donkey anti-rabbit (cat. 925-32213).

### Expression constructs

For in vitro transcription/translation, full length human VPS11, VPS16, VPS18, VPS33A, VPS39 and VPS41, with restriction sites silently mutated as described in Graham et al 2013 (18), and VPS8, isolated from HeLa cell cDNA, were cloned into pF3A WG (BYDV) (Promega) with N-terminal myc tags. Full length and C-terminal truncation constructs of VPS8, VPS18 and VPS41 had an EGFP tag added to the C terminus by cloning into pEGFP-N1 (Clontech). The RING domain and other N-terminal truncation constructs had the EGFP tag added to the N terminus by cloning into pEGFP-C1 (Clontech). For expression in E. coli, the RING domains of human VPS18 (residues 842–973), VPS41 (residues 778–854) and VPS8 (short: residues 1246–1320; long, including C-terminal ‘disordered’ region: residues 1246–1426) were cloned into pOPT3G (18), encoding an N-terminal GST tag and human rhinovirus 3C protease sequence. The RING domain of VPS18 was cloned into pOPTnH (48), encoding a C-terminal LysHis6 tag. For yeast two-hybrid analysis, the putative RING domains of VPS18 and VPS41, truncated VPS18 RING domain constructs (ΔC1 and ΔC2, spanning residues 842–957 and 842–961, respectively), and the zinc finger domain of VPS39 (residues 840–875), were cloned into pGBKT7 (Takara).

### Yeast two-hybrid screening

Yeast two-hybrid experiments were performed using the Matchmaker Gold yeast two-hybrid system (Takara) as per the manufacturer’s instructions. VPS18 RING ΔC2, VPS41 RING and the VPS39 zinc finger were used as baits against the Mate and Plate Universal Human (Normalised) library. For each, 6–18 million clones were screened and putative interaction partners were identified by PCR of positive colonies. Selected putative interaction partners were cloned with an N-terminal myc tag into pF3A WG (BYDV), expressed using cell-free transcription/translation in wheat germ lysate and probed for their ability to bind the purified bait protein (see below), but for none could a positive interaction be confirmed.

### Protein expression and purification

In vitro protein expression was performed using TNT SP6 High Yield Wheat Germ reaction mix (Promega) as per the manufacturer’s instructions. GST-tagged proteins were expressed in E. coli BL21(DE3) pLysS (ThermoFisher) and His-tagged VPS18 was expressed in E. coli Rosetta2(DE3) pLysS (Novagen). Bacteria were grown in 2×TY medium to an A_600_ of 0.8–1.2 at 37°C, cooled to 22°C, and protein expression was induced by the addition of 0.2 mM isopropyl β-d-thiogalactopyranoside. After 16 h, cells were harvested by centrifugation at 5000×g for 15 min and the pellet was stored at −20°C until required.

For GST-tagged proteins, cells were thawed and resuspended in 20 mM Tris (pH 7.5), 300 mM NaCl, 1.4 mM β-mercaptoethanol, 0.05% TWEEN 20, supplemented with 400 units of bovine DNase I (Sigma-Aldrich) and 200 μL of EDTA-free protease inhibitor mixture (Sigma-Aldrich) per 2 L of cell culture. Cells were lysed at 24 kpsi using a TS series cell disruptor (Constant Systems) and lysates cleared by centrifugation at 40,000×g for 30 min at 4°C. Cleared lysate was incubated with glutathione sepharose 4B (GE Healthcare) for 1 h at 4°C, the beads were washed with 20 mM Tris pH 7.5, 300 mM NaCl, 1 mM DTT, and bound protein was eluted in wash buffer supplemented with 25 mM reduced glutathione. His-tagged VPS18 RING was purified by following the same general procedure using Ni^2+^-NTA immobilisation resin (Qiagen) and the following buffers: *lysis buffer*, 20 mM Tris (pH 7.5), 20 mM imidazole, 500 mM NaCl, 1.4 mM β-mercaptoethanol, 0.05% TWEEN 20 supplemented with 400 U bovine DNase I and 200 μl EDTA-free protease inhibitor cocktail per 2 L of bacterial culture; *wash buffer*, 20 mM Tris (pH 7.5), 20 mM imidazole, 500 mM NaCl; *elution buffer*, 20 mM Tris (pH 7.5), 250 mM imidazole, 500 mM NaCl. All affinity-purified proteins were injected onto 16/600 Superdex 200 (GST-tagged proteins) or 75 (VPS18 RING H_6_) size exclusion chromatography (SEC) columns (GE Healthcare) equilibrated in 20 mM Tris (pH 7.5), 200 mM NaCl, 1 mM DTT (SEC buffer). The GST tag of GST-VPS41 RING was removed by incubation overnight at 4°C with 50–100 μg of human rhinovirus 3C protease in SEC buffer supplemented with 0.5 mM EDTA and fresh DTT (final concentration 2 mM). Following capture of GST and uncleaved GST-VPS41 RING using glutathione sepharose 4B, VPS41 RING was further purified by SEC using a 16/600 Superdex 75 column as described above. All purified proteins were concentrated, snap-frozen in liquid nitrogen, and stored at −80 °C until required.

### GST pulldown experiments

Affinity capture (GST pulldown) experiments were performed in 96-well flat-bottomed plates (Greiner) using magnetic glutathione beads (Thermo Scientific). For capture of prey proteins produced by cell-free expression in wheat germ lysate, 0.5 nmol of purified GST-tagged bait protein was incubated with magnetic glutathione beads at 4°C for 10–20 min. The beads were washed three times with wash buffer (20 mM Tris, 200 mM NaCl, 0.1% NP-40, 1 mM DTT, 1 mM EDTA), incubated with wheat germ cell-free expression reactions at 4°C for 60 min, and then washed four times in wash buffer. Bound protein was eluted using wash buffer supplemented with 50 mM reduced glutathione and then boiled in SDS-PAGE loading buffer for SDS-PAGE and immunoblotting. For capture of purified prey proteins expressed in E. coli, 1.5 nmol of GST-tagged bait protein was incubated with magnetic glutathione beads for 15 min at 4°C and then washed three times with 20 mM Tris, 200 mM NaCl, 0.1% NP-40, 1 mM DTT. The bait-loaded beads were incubated with 1.5 nmol of purified prey protein at 4°C for 60 min, washed four times, and bound protein was eluted using wash buffer supplemented with 50 mM reduced glutathione before boiling in SDS-PAGE loading buffer for SDS-PAGE analysis.

### Multi-angle light scattering

Multiangle light scattering (MALS) experiments were performed at room temperature by inline measurement of static light scattering (DAWN 8+, Wyatt Technology), differential refractive index (Optilab T-rEX, Wyatt Technology), and 280 nm absorbance (Agilent 1260 UV, Agilent Technologies) following SEC at a flow rate of 0.5 ml/min. Samples (100 μL) were injected onto an analytical Superdex 75 10/300 gel filtration column (GE Healthcare) equilibrated in 20 mM Tris (pH 7.5), 150 mM NaCl, 0.5 mM EDTA. VPS18 RING H_6_ was injected at 1.16 mg/mL, VPS41 RING at 1.33 mg/mL, and the VPS18:VPS41 complex was formed by incubating 0.48 mg/mL VPS41 RING with 0.68 mg/mL VPS18 RING H_6_ (yielding a 1.0:0.75 molar ratio) on ice for 15 min before injection. Molar masses were calculated using ASTRA 6 (Wyatt Technology).

### Cell culture and transfection

HEK293T cells (ATCC #CRL-3216, mycoplasma-free) were grown in Dulbecco’s Modified Eagle Medium with high glucose (Sigma, cat. D6546), supplemented with 10% (v/v) heat-inactivated foetal calf serum, 2 mM L-glutamine, 100 IU/mL penicillin and 100 mg/mL streptomycin, in a humidified 5% CO_2_ atmosphere at 37°C. Cells were transfected with TransIT-LT1 (Mirus), following the manufacturer’s instructions.

### Immunoprecipitation and immunoblotting

Transfected HEK293T cells were harvested and lysed in 10 mM Tris-HCl pH 7.5, 150 mM NaCl, 0.5 mM EDTA, 0.5% NP-40, and EDTA-free protease inhibitor cocktail (Sigma). Protein concentration in the resulting lysates was quantified by BCA assay (Thermo Scientific), and protein concentration was equalised before immunoprecipitation with GFP-TRAP (ChromoTek) beads according to the manufacturer’s protocol.

Protein samples were boiled in SDS-PAGE loading buffer for 5 minutes before separation on a 10% SDS-PAGE gel and transferring to nitrocellulose membrane (Protran) using the Mini-PROTEAN system (BioRad). After blocking in PBS with 5% (w/v) non-fat milk powder, membranes were incubated with primary antibody overnight at 4°C, then secondary antibody for 1 hour at room temperature. Dried blots were visualised on an Odyssey infrared scanner (LI-COR)

## ACKNOWLEDGEMENTS

We thank Janet Deane for assistance with MALS experiments, Sally Gray for assistance with yeast two-hybrid experiments, Lena Wartosch and Paul Luzio for reagents and helpful comments, and Yvonne Hackmann for HeLa cell cDNA.

## AUTHOR CONTRIBUTIONS

SCG conceived the project; MRH, EJS, EE and SCG designed and performed experiments; MRH, EJS and SCG analysed the data; MRH and SCG wrote the paper; all authors revised and approved the final manuscript.

---

**Supplementary figure 1:**
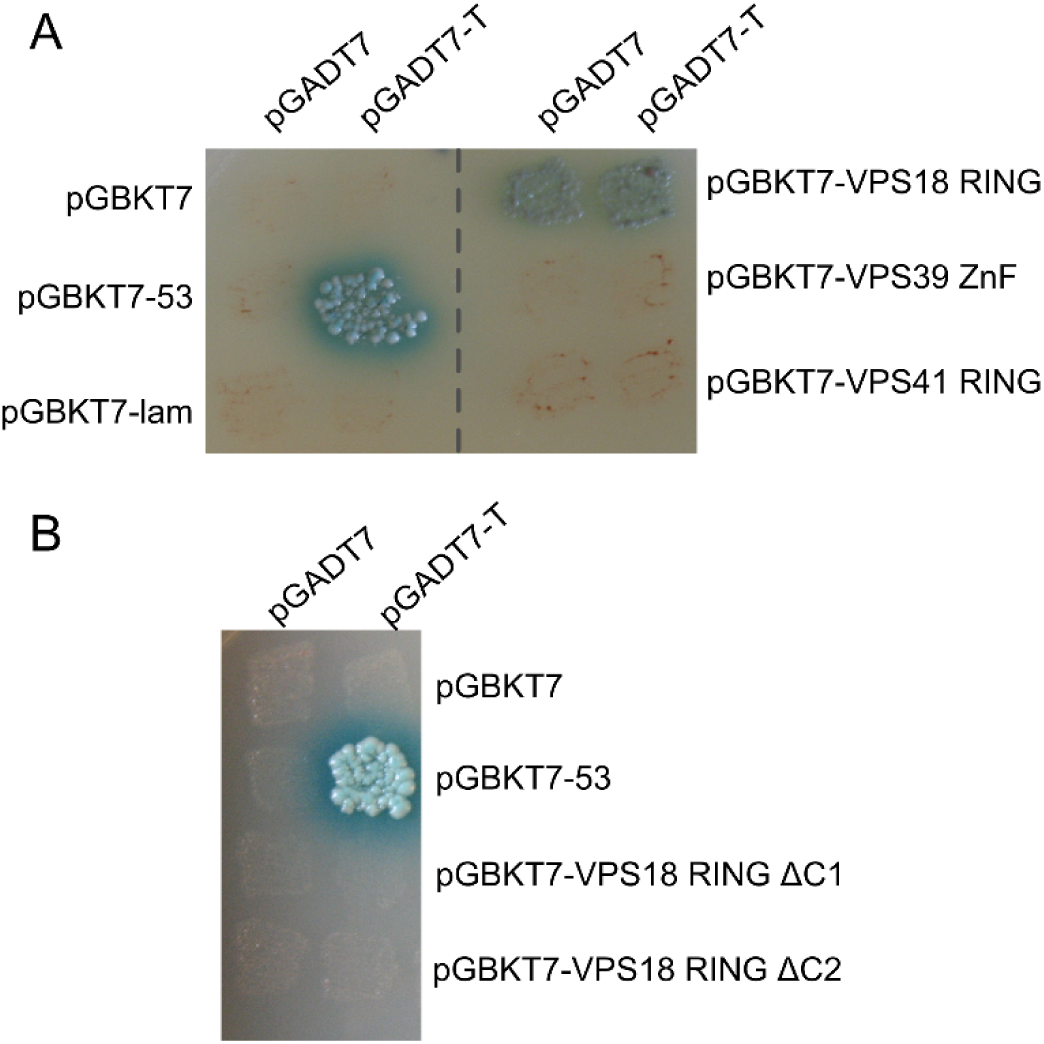
Auto-activation testing of HOPS RING domains for yeast two-hybrid experiments. (A) Yeast strain Y2H Gold was transformed with a Gal4 DNA binding domain fused to VPS18 and VPS41 RING domains (pGBKT7-VPS18 RING and pGBKT7-VPS41 RING), to the VPS39 zinc finger domain (pGBKT7-VPS39 ZnF), to a positive control (human p53; pGBKT7-53), to a negative control (human lamin C; pGBKT7-lam), or in isolation (pGBKT7). Strain Y187 was transformed with the Gal4 activation domain in isolation (pGADT7) or fused to the SV40 large T antigen (pGADT7-T). Yeast were mated and then grown on -Leu -Trp -Ade -His nutritional selection media supplemented with X-α-Gal and aureobasidin A, growth of blue colonies indicating a positive interaction. (B) Two truncated VPS18 RING domain constructs where the putative C-terminal helix had been removed (VPS18 RING ΔC1 and ΔC2, spanning residues 842–957 and 842–961, respectively) were tested for auto-activation as described in (A).

**Supplementary figure 2:**
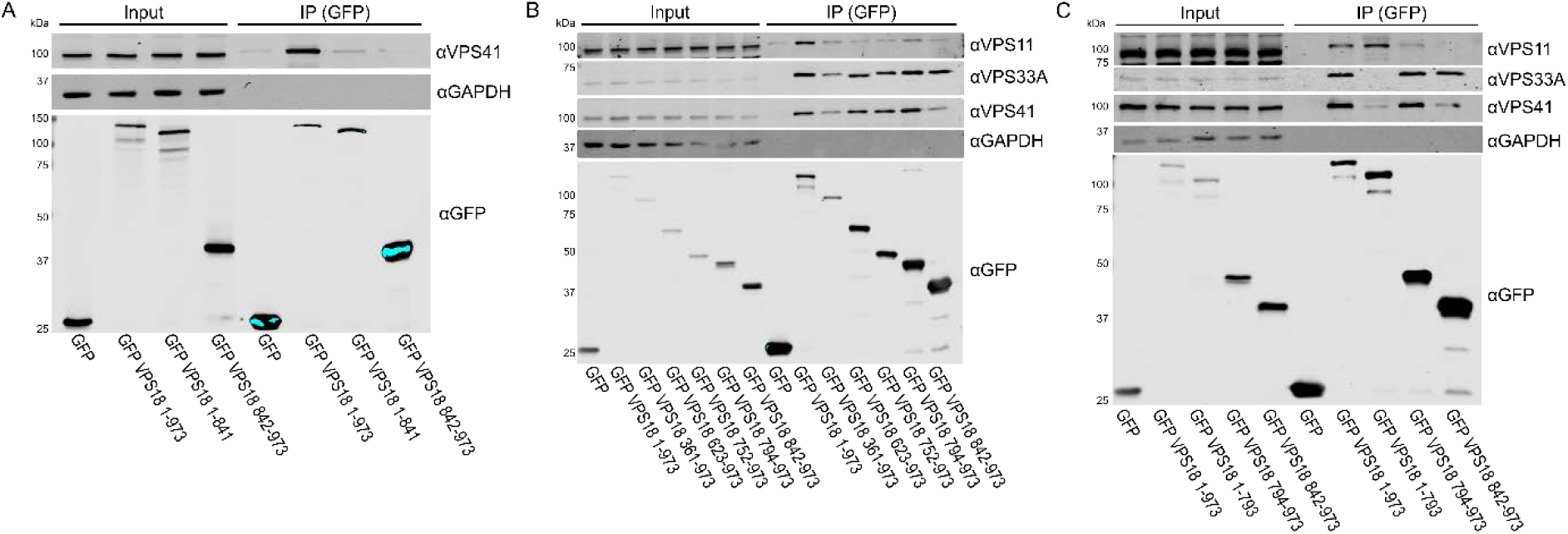
Co-immunoprecipitations of various VPS18 truncation constructs expressed in HEK293T cells. (A) The VPS18 (1-841) construct retains some interaction with endogenous VPS41. (B) A series of N-terminal truncations of VPS18 shows that extended RING domain constructs are more efficient at co-immunoprecipitating endogenous VPS41 than the RING domain alone (842-973). (C) VPS18 (1-793) co-immunoprecipitates endogenous VPS11 with an efficiency equal to full length VPS18 (1-973).

**Supplementary data: Proteins identified as putative HOPS RING domain interaction partners by yeast two-hybrid.** See Excel file.

## REFERENCES

1. Baker RW, Jeffrey PD, Zick M, Phillips BP, Wickner WT, Hughson FM. A direct role for the Sec1/Munc18-family protein Vps33 as a template for SNARE assembly. Science. 2015;349(6252):1111–4.

2. Baker RW, Hughson FM. Chaperoning SNARE assembly and disassembly. Nature reviews Molecular cell biology. 2016;17(8):465–79.

3. Balderhaar, HJk, Ungermann C. CORVET and HOPS tethering complexes – coordinators of endosome and lysosome fusion. Journal of Cell Science. 2013;126(6):1307–16.

4. Spang A. Membrane tethering complexes in the endosomal system. Frontiers in cell and developmental biology. 2016;4.

5. Perini ED, Schaefer R, Stöter M, Kalaidzidis Y, Zerial M. Mammalian CORVET is required for fusion and conversion of distinct early endosome subpopulations. Traffic. 2014;15(12):1366–89.

6. Wartosch L, Günesdogan U, Graham SC, Luzio JP. Recruitment of VPS33A to HOPS by VPS16 Is Required for Lysosome Fusion with Endosomes and Autophagosomes. Traffic. 2015;16(7):727–42.

7. Jiang P, Nishimura T, Sakamaki Y, Itakura E, Hatta T, Natsume T, et al. The HOPS complex mediates autophagosome-lysosome fusion through interaction with syntaxin 17. Mol Biol Cell. 2014;25(8):1327–37.

8. Liang C, Lee JS, Inn KS, Gack MU, Li Q, Roberts EA, et al. Beclin1-binding UVRAG targets the class C Vps complex to coordinate autophagosome maturation and endocytic trafficking. Nature cell biology. 2008;10(7):776–87.

9. Takats S, Pircs K, Nagy P, Varga A, Karpati M, Hegedus K, et al. Interaction of the HOPS complex with Syntaxin 17 mediates autophagosome clearance in Drosophila. Mol Biol Cell. 2014;25(8):1338–54.

10. Klinger CM, Klute MJ, Dacks JB. Comparative genomic analysis of multi-subunit tethering complexes demonstrates an ancient pan-eukaryotic complement and sculpting in Apicomplexa. PloS one. 2013;8(9):e76278.

11. van der Kant R, Jonker CTH, Wijdeven RH, Bakker J, Janssen L, Klumperman J, et al. Characterization of the Mammalian CORVET and HOPS Complexes and Their Modular Restructuring for Endosome Specificity. Journal of Biological Chemistry. 2015;290(51):30280–90.

12. Lachmann J, Glaubke E, Moore PS, Ungermann C. The Vps39-like TRAP1 is an effector of Rab5 and likely the missing Vps3 subunit of human CORVET. Cellular logistics. 2014;4(4):e970840.

13. Rink J, Ghigo E, Kalaidzidis Y, Zerial M. Rab conversion as a mechanism of progression from early to late endosomes. Cell. 2005;122(5):735–49.

14. van der Kant R, Fish A, Janssen L, Janssen H, Krom S, Ho N, et al. Late endosomal transport and tethering are coupled processes controlled by RILP and the cholesterol sensor ORP1L. J Cell Sci. 2013;126(15):3462–74.

15. Bröcker C, Kuhlee A, Gatsogiannis C, kleine Balderhaar HJ, Hönscher C, Engelbrecht-Vandré S, et al. Molecular architecture of the multisubunit homotypic fusion and vacuole protein sorting (HOPS) tethering complex. Proceedings of the National Academy of Sciences. 2012;109(6):1991–6.

16. Chou H-T, Dukovski D, Chambers MG, Reinisch KM, Walz T. CATCHR, HOPS and CORVET tethering complexes share a similar architecture. Nat Struct Mol Biol. 2016;23(8):761–3.

17. Gissen P, Johnson CA, Gentle D, Hurst LD, Doherty AJ, O’Kane CJ, et al. Comparative evolutionary analysis of VPS33 homologues: genetic and functional insights. Human Molecular Genetics. 2005;14(10):1261–70.

18. Graham SC, Wartosch L, Gray SR, Scourfield EJ, Deane JE, Luzio JP, et al. Structural basis of Vps33A recruitment to the human HOPS complex by Vps16. Proceedings of the National Academy of Sciences. 2013;110(33):13345–50.

19. Suzuki T, Oiso N, Gautam R, Novak EK, Panthier J-J, Suprabha P, et al. The mouse organellar biogenesis mutant buff results from a mutation in Vps33a, a homologue of yeast vps33 and Drosophila carnation. Proceedings of the National Academy of Sciences. 2003;100(3):1146–50.

20. Kondo H, Maksimova N, Otomo T, Kato H, Imai A, Asano Y, et al. Mutation in VPS33A affects metabolism of glycosaminoglycans: a new type of mucopolysaccharidosis with severe systemic symptoms. Human Molecular Genetics. 2017;26(1):173–83.

21. Gissen P, Johnson CA, Morgan NV, Stapelbroek JM, Forshew T, Cooper WN, et al. Mutations in VPS33B, encoding a regulator of SNARE-dependent membrane fusion, cause arthrogryposis–renal dysfunction–cholestasis (ARC) syndrome. Nature genetics. 2004;36(4):400–4.

22. Gruber R, Rogerson C, Windpassinger C, Banushi B, Straatman-Iwanowska A, Hanley J, et al. Autosomal Recessive Keratoderma-Ichthyosis-Deafness (ARKID) Syndrome Is Caused by VPS33B Mutations Affecting Rab Protein Interaction and Collagen Modification. Journal of Investigative Dermatology. 2017;137(4):845–54.

23. Urban D, Li L, Christensen H, Pluthero FG, Chen SZ, Puhacz M, et al. The VPS33B-binding protein VPS16B is required in megakaryocyte and platelet α-granule biogenesis. Blood. 2012;120(25):5032–40.

24. Brett CL, Plemel RL, Lobingier BT, Vignali M, Fields S, Merz AJ. Efficient termination of vacuolar Rab GTPase signaling requires coordinated action by a GAP and a protein kinase. J Cell Biol. 2008;182(6):1141–51.

25. Khatter D, Raina VB, Dwivedi D, Sindhwani A, Bahl S, Sharma M. The small GTPase Arl8b regulates assembly of the mammalian HOPS complex on lysosomes. Journal of Cell Science. 2015;128(9):1746–61.

26. Garg S, Sharma M, Ung C, Tuli A, Barral Duarte C, Hava David L, et al. Lysosomal Trafficking, Antigen Presentation, and Microbial Killing Are Controlled by the Arf-like GTPase Arl8b. Immunity. 2011;35(2):182–93.

27. Behrmann H, Lürick A, Kuhlee A, Balderhaar HK, Bröcker C, Kümmel D, et al. Structural Identification of the Vps18 β-Propeller Reveals a Critical Role in the HOPS Complex Stability and Function. Journal of Biological Chemistry. 2014;289(48):33503–12.

28. Rieder SE, Emr SD. A Novel RING Finger Protein Complex Essential for a Late Step in Protein Transport to the Yeast Vacuole. Molecular Biology of the Cell. 1997;8(11):2307–27.

29. Nickerson DP, Brett CL, Merz AJ. Vps-C complexes: gatekeepers of endolysosomal traffic. Current Opinion in Cell Biology. 2009;21(4):543–51.

30. Robinson JS, Graham TR, Emr SD. A putative zinc finger protein, Saccharomyces cerevisiae Vps18p, affects late Golgi functions required for vacuolar protein sorting and efficient alpha-factor prohormone maturation. Molecular and Cellular Biology. 1991;11(12):5813–24.

31. Radisky DC, Snyder WB, Emr SD, Kaplan J. Characterization of VPS41, a gene required for vacuolar trafficking and high-affinity iron transport in yeast. Proceedings of the National Academy of Sciences of the United States of America. 1997;94(11):5662–6.

32. Huizing M, Didier A, Walenta J, Anikster Y, Gahl WA, Krämer H. Molecular cloning and characterization of human VPS18, VPS 11, VPS16, and VPS33. Gene. 2001;264(2):241–7.

33. Guo Z, Johnston W, Kovtun O, Mureev S, Bröcker C, Ungermann C, et al. Subunit organisation of in vitro reconstituted HOPS and CORVET multisubunit membrane tethering complexes. PloS one. 2013;8(12):e81534.

34. Borden KLB, Freemont PS. The RING finger domain: a recent example of a sequence—structure family. Current Opinion in Structural Biology. 1996;6(3):395–401.

35. Saurin AJ, Borden KL, Boddy MN, Freemont PS. Does this have a familiar RING? Trends in biochemical sciences. 1996;21(6):208–14.

36. Borden KL. RING fingers and B-boxes: zinc-binding protein-protein interaction domains. Biochemistry and Cell Biology. 1998;76(2-3):351–8.

37. Mizuno K, Kitamura A, Sasaki T. Rabring7, a Novel Rab7 Target Protein with a RING Finger Motif. Molecular Biology of the Cell. 2003;14(9):3741–52.

38. Kentsis A, Gordon RE, Borden KL. Self-assembly properties of a model RING domain. Proceedings of the National Academy of Sciences of the United States of America. 2002;99(2):667–72.

39. Petersen B, Petersen TN, Andersen P, Nielsen M, Lundegaard C. A generic method for assignment of reliability scores applied to solvent accessibility predictions. BMC structural biology. 2009;9(1):51.

40. Alva V, Nam S-Z, Söding J, Lupas AN. The MPI bioinformatics Toolkit as an integrative platform for advanced protein sequence and structure analysis. Nucleic acids research. 2016;44(W1):W410–W5.

41. Capili AD, Schultz DC, Rauscher FJ, Borden KLB. Solution structure of the PHD domain from the KAP-1 corepressor: structural determinants for PHD, RING and LIM zinc-binding domains. The EMBO Journal. 2001;20(1-2):165–77.

42. Yang F, Hsu P, Lee SD, Yang W, Hoskinson D, Xu W, et al. The C terminus of Pcf11 forms a novel zinc-finger structure that plays an essential role in mRNA 3’-end processing. RNA (New York, NY). 2017;23(1):98–107.

43. McVey Ward D, Radisky D, Scullion MA, Tuttle MS, Vaughn M, Kaplan J. hVPS41 Is Expressed in Multiple Isoforms and Can Associate with Vesicles through a RING-H2 Finger Motif. Experimental Cell Research. 2001;267(1):126–34.

44. Mackay JP, Sunde M, Lowry JA, Crossley M, Matthews JM. Protein interactions: is seeing believing? Trends in Biochemical Sciences. 2007;32(12):530–1.

45. Epp N, Ungermann C. The N-terminal domains of Vps3 and Vps8 are critical for localization and function of the CORVET tethering complex on endosomes. PLoS One. 2013;8(6):e67307.

46. Aravind L, Iyer L, Koonin EV. Scores of RINGS but no PHDs in ubiquitin signaling. Cell Cycle. 2003;2(2):123–6.

47. Dodd RB, Allen MD, Brown SE, Sanderson CM, Duncan LM, Lehner PJ, et al. Solution Structure of the Kaposi’s Sarcoma-associated Herpesvirus K3 N-terminal Domain Reveals a Novel E2-binding C4HC3-type RING Domain. Journal of Biological Chemistry. 2004;279(51):53840–7.

48. Neidel S, de Motes CM, Mansur DS, Strnadova P, Smith GL, Graham SC. Vaccinia virus protein A49 is an unexpected member of the B-cell Lymphoma (Bcl)-2 protein family. Journal of Biological Chemistry. 2015;290(10):5991–6002.

